# Rational synthesis of total damage during vitrification: modelling and experimental validation of osmotic, temperature, and cytotoxic damage in sea urchin (*Paracentrotus lividus)* oocytes

**DOI:** 10.1101/2022.07.08.499206

**Authors:** Dominic J Olver, Pablo Heres, Estefania Paredes, James D Benson

## Abstract

Sea urchin (*Paracentrotus lividus*) oocytes are an important species for aquaculture and as a model species for multiple scientific fields. Despite their importance, methods of cryopreserved biobanking of oocytes are currently not possible. Optimized cryoprotectant loading may enable vitrification methods of cryopreservation and thus long-term storage of oocytes. Determining an optimized protocol requires membrane characteristics and models of damage associated with the vitrification loading protocol, namely osmotic, temperature, and cytotoxic damage. We present and experimentally evaluated state-of-the-art models alongside our novel models. We experimentally verify the damage models throughout time at difference treatment intensities. Osmotic damage experiments consisted of hypertonic solutions composed of seawater supplemented with NaCl or sucrose and hypotonic solutions composed of seawater diluted with deionized water. Treatment times ranged from 2 to 30 minutes. To test temperature damage (in particular chill injury), oocytes were exposed to 1.7 °C, 10 °C, and 20 °C (control) for exposure times ranging from 2 to 90 minutes. Cytotoxicity was investigated by exposing oocytes to solutions of Me_2_SO for exposure times ranging from 2 to 30 minutes. We identify appropriate models and use these to search for an optimal loading protocol, namely the time dependent osmotic damage model (for osmotic damage), the temperature dependent model (for temperature damage), and the external molality Arrhenius power model (for cytotoxicity). We combined these models to estimate total damage during a cryopreservation loading protocol and performed a exhaustive grid search for optimal loading for a given goal intracellular cryoprotectant concentration. Given our fitted models, we find sea urchin oocytes can only be loaded to 0.13 Me_2_SO v/v with a 50% survival, For reference, levels for vitrification are approximately 0.45 v/v. Our synthesis of damages is the first of its kind, and enables a fundamentally novel approach to modelling survival for cells in general.

## Introduction

Sea urchins are a long standing model system in developmental biology, embryology, and aquaculture (1–4). Successful deployment of such biobanking techniques will enable efficient storage and transfer of gametes aiding scientific and aquacultural endeavors (5–9). However, for sea urchins, cryopreservation methods of preserving female gametes or embryos of sea urchins are very difficult and often unsuccessful with the highest recovery rate of 10% for fertilized eggs (10) or non−existent for oocytes (9, 11). During cryopreservation, cells may undergo volumetric change, exposure to high concentrations of cytotoxic solutions, and exposure to cold temperatures, all potentially leading to loss of cell viability (12–14). Surprisingly, little is known about the mechanisms behind osmotic damage, cytotoxicity, and chill injury in the context of cryobiology, and by extension mathematical models to help predict (and therefore avoid) damages are not wildly used or well verified. Development of a comprehensive damage model that includes these major sources of damage during cryopreservation will enable a rational approach to optimized cryopreservation protocols along with illuminating mechanisms of damage.

Vitrification is a common procedure for oocyte cryopreservation in part due to the avoidance of ice formation (7, 13). However, vitrification calls for solutions including cryoprotective agents (CPAs) that are 40% to 50% of the solution volume. In these solutions (ranging from 3 to 9 osmol/kg of CPAs), cells may experience high osmotic pressure gradients during the vitrification protocol. The osmotic gradient forces water out of the cell during loading while membrane permeable CPA diffuses into the cell, resulting in the initial cell volume loss (water leaving the cell) and then subsequent swelling (CPA + water entering the cell). The so called two−parameter (2P) model is a useful model describing the flux of water and CPA across the membrane and is commonly expanded to include temperature dependence (14, 15). This 2P model requires information about non−osmotically active volume of the cell and assumes the cell to be an “ideal osmometer”. The Boyle van ‘t Hoff relation provides non-osmotically active volume and determines if the cell is an “ideal osmometer” (16, 17).

With large osmotic gradients comes large changes in cell volume. Classically, cells are thought to have osmotic tolerance limits (OTLs), that is, the limit that a cell can shrink or swell to without dying from mechanical stress (18, 19). Additionally, cytotoxic damage may result from the presence of CPA in the cell (20, 21). The longer the CPA is inside the cell, the more cytotoxic damage accumulates, ultimately resulting in irreversible damage and cell death. An optimized vitrification protocol is one that achieves the proper intracellular CPA concentration for successful vitrification while minimizing cytotoxicity and avoiding osmotic damage (14, 22).

Zawlodzka and Takamatsu (23) found that osmotic damage is related to not just maximum osmotic challenge but also the time of exposure. However, time dependent osmotic damage is poorly understood; while there is no direct evidence for the mechanism of osmotic damage, the Zawlodzka and Takamatsu hypothesize there is ultrastructural alteration to the cell as well as reduction of total membrane during shrinkage as described by membrane regulation hypothesis (24). Such changes to the cell membrane along with alterations to the cytoskeleton may result in catastrophic damage to the cell upon return to isotonic volume. Since these ultrastructural alterations, membrane adaption and cytoskeletal changes are thought to be time dependent, it follows that the associated damage is also time dependent. Indeed, osmotic damage has been found to correlate to the accumulation of volumetric deviance throughout time (25, 26).

Experimentally, mitigation of osmotic damage may be obtained by alterations to the cytoskeleton and cell membrane (27–29). Cytochalasin-B aids in vitrification for porcine and buffalo oocytes (27, 29); this molecule degrades the cytoskeleton and prevents reformation during the shrink−swell cycle associated with the vitrification process. Further, it is found that addition of cholesterol also aids vitrification of cells (28). Possible reasons include increased cell membrane fluidity and can regulate the connectivity between the cell membrane and the cytoskeleton via Phosphatidylinositol 4,5-bisphosphate (PIP_2_) managed connections (30–32). Cells with added cholesterol can exhibit similar membrane dynamics as cells with denatured cytoskeletons (33). It is possible that the benefits from addition of cytochalasin-B or cholesterol are achieved by retarding time dependent reformation of the cytoskeleton during CPA equilibrium. If volumetric damage is proportional to the accumulated change in volume throughout time, then we expect a relationship dependent on time of exposure and osmolality that may be mathematically modeled.

Cytotoxicity of CPAs is a key limitation of cryopreservation and the major limitation for vitrification protocols (34). Cytotoxic damage accumulates during both the addition and removal of CPAs and is dependent on CPA concentration, time, and temperature (35–38). The damage due to CPA toxicity is complex and poorly understood. Despite the complexity, cytotoxicity can still be modelled (20–22,36). The model presented by Benson and colleagues (20) assumes cytotoxicity is accumulated throughout time and dependent on CPA concentration in the cell. It is theorized that at lower temperatures cytotoxicity builds up slower, however flux of CPA across the membrane is also slower (36). Therefore, there is a trade-off between the amount of time required to load/unload a cell with CPA and the cytotoxicity accumulated throughout that time. This trade-off is a function of CPA concentration and environmental temperature. However, some cells are sensitive to low temperatures (even in the absence of ice) (39).

Chill injury is injury to cells at low temperatures in the absence of ice (e.g. 0 to 10 °C) (39–41). Chill injury is documented in multiple fields but is not well understood (39–41). Both historic and current models of chill injury use simple sigmoidal fits relating time, and temperature with survival (39,42,43). Importantly, the time frame for these models is in terms of hours or days-much longer than cryobiological protocols which are usually on the order of minutes to tens of minutes for oocytes. In cryobiology, chill injury is hypothesized to be primarily caused by meta−structural changes in the cell membrane during cool temperatures (40). During cooling, lipids undergo a phase transition and upon return to physiological temperatures the lipid membrane would become compromised and fail (40). However, other possible mechanisms of chill injury include ionic instability during cooling (44–46). Despite these hypotheses, there has been no conclusive evidence of a known mechanism of chill injury nor a physiologically derived mathematical model capable of predicting chill injury on the time period of tens of minutes. Indeed, to our knowledge, there has been no consideration of chill injury in optimizing vitrification or slow cooling protocols. Despite the lack of a mathematical model, equilibrating cells with CPA at low temperatures may be a good method for avoiding high toxicity levels (22, 36). Indeed, such a method is required for liquidous tracking (47). However, if too many cells die from chill injury, then this method may not be viable. Therefore, a model that predicts chill injury on the time scale of minutes to tens of minutes is necessary for obtaining an optimized vitrification protocol across both time and temperature.

The object of this paper is to present and experimentally verify novel mathematical models of cell damage during cryopreservation protocols for sea urchin (*Paracentrotus lividus*) oocytes. First, we present and develop multiple damage model types: cytotoxic damage, osmotic damage, and temperature damage (chill injury). Next, we determine cellular characteristics (Boyle van ‘t Hoff relation and fit 2P model). Using these characteristics, we experimentally verify damage models for osmotic damage, temperature damage, and cytotoxicity. We find that osmotic damage is time dependent, chill injury is temperature dependent but not time dependent, and cytotoxicity is best predicted by tracking CPA concentration at the membrane. Finally, we combine the verified models and perform an exhaustive grid search of an optimized continuous loading protocol for multiple goal intracellular CPA concentrations and find that sea urchin oocytes can only be loaded to approximately 15% CPA volume per solution volume (v/v).

## Materials and Methods

### Modeling

#### Boyle van ‘t Hoff relation

The Boyle van ‘t Hoff (BvH) relation describes the relationship between equilibrated relative cell volume, *v*, with respect to osmotic pressure, *Π*, where ν = *V/V*_iso_, *V*, is cell volume and *V*_iso_ is cell volume at isotonic pressure, *Π*_iso_ (16,17,48). Noting the cell is composed of osmotically active volume and osmotically inactive volume (solids), then relative volume may be written in terms of normalized osmotically active fraction, *w*_iso_, and osmotically inactive fraction, *b*, of the cell (i.e. *v*=*w*_iso_+*b*). The BvH relation takes the form,

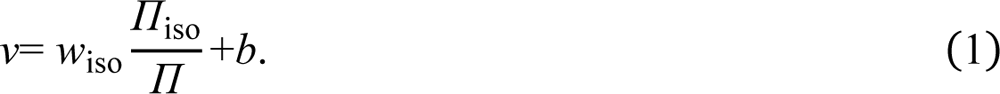

The fitted, *w*_iso_, and, *b*, parameters inform other models such as the two parameter (2P) model to enable time dependent predictions of volumetric change with respect to osmotic conditions (49, 50). We note that Eq. 1 does not necessarily need to be normalized with respect to the isotonic point but is done here for simplicity and for the direct extraction of *w*_iso_.

#### Two Parameter (2P) Model

The 2P model describes the flux of water and membrane permeable CPA across the cell membrane at a given moment in time, *t*. Originally, Jacobs (51) presented a model of the mass transport of water and permeable solutes across the membrane in terms of chemical potentials. For water, *W*, this process may be written in terms of an osmotic pressure gradient, whereas mols of CPA, *S*, it is written in terms of a concentration gradient (14). The flux of water and CPA across the cell membrane may be written in terms of a system of ordinary differential equations in the form,

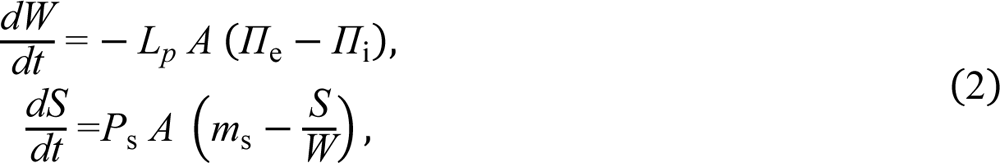

where *A* is the surface area of the cell (often assumed constant), *m*_s_ is the extracellular molality of membrane permeable CPA, *L*_p_ is the hydraulic conductivity, and *P*_s_ is the CPA permeability constant. The external osmotic pressure may be written as *Π*_e_=*RT*(*π*_n_+*π*_s_) and the intracellular osmotic pressure may be written as *Π*_i_=*RT*(*N*_i_+*S*)/*W*, where *R* is the gas constant, *T* is temperature, *π_n_* is the external osmolality of non−permeable ion/osmolytes, *π*_s_ is extracellular osmolality of permeable CPA, and *N*_i_ is the osmols of non−permeable internal ions/osmolytes. Typically, the 2P model is fit to experimental data for cell volume throughout CPA equilibration. The hydraulic conductivity, *L*_p_, and CPA permeability constant, *P*_s_, are both assumed to follow an Arrhenius equation, taking the form,

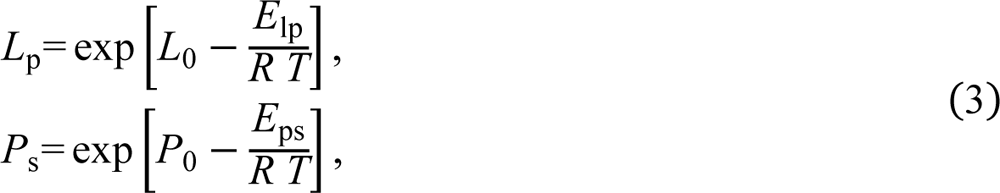

where *L*_0_, *P*_0_, *E*_lp_, and *E*_ps_ are fitted constants. These equations can be fit linearly by taking the log of *L*_p_ and *P*_s_ values at different temperatures and plotting them with respect to inverse temperature *T*^−1^ (in kelvin), creating a linear relation (e.g. log[*L*_p_]*=L*_0_-*E*_lp_×*R*^−1^×*T* ^−1^).

#### Cytotoxicity model

A population death rate model provides a first order rate equation to model cell death after exposure to Me_2_SO (21,36–38). This population death rate is presented as an ordinary differential equation and takes the form,

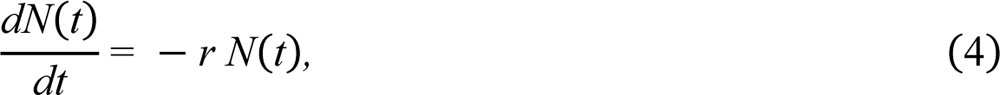

where *dN*(*t*) is the rate of change in surviving population size at some time *t*, and *r* is the decay rate. Solving Eq. 4 with respect to some time provides the form,

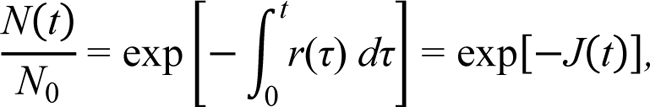

where *N*_0_ is initial population, *τ* is all time points between the start and the time *t*, *J*(*t*) is the accumulated damage function for the population up to some time *t* and represented as the integration of the decay rate.

Benson and colleagues (21) used this approximation to define cytotoxicity, where decay rate is proportional to a power function of intracellular CPA molality *m*_i_. The relative amount of population surviving at the final time is equal to the damage accrued *J*_tox_ throughout time such that,

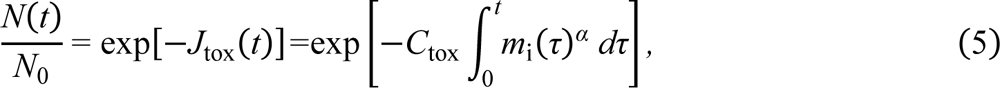

where *C*_tox_ and *α* are fitted constants, *m*_i_(*t*)*=S*(*t*)*/W*(*t*), *S*(*t*) and *W*(*t*) are intracellular moles of CPA and mass of water at some time *t* respectively. Davidson and colleagues (36) extend Eq. 5 across temperature by allowing *C*_tox_ be a function of temperature governed by the Arrhenius equation (i.e. *C*tox(*T*) = exp[*C*0−*E*tox/*RT*]), taking the form,

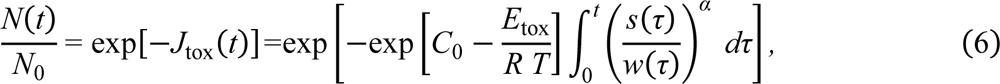

where *C*_0_ and *E*_tox_ are fitted parameters. Eq. 6 will be referred to as the Intracellular Molality Arrhenius (IMA) model. The power parameter, *α*, may be interpreted as a sensitivity threshold parameter, such that highly sensitive cells have a high *α* value. In practicality, α controls how closely bunched cell survival curves are with respect to intracellular molality as a function of time, while the proportionality constant *C*_tox_ governs the slope of cell survival as a function of time. If cell sensitivity changes with respect to temperature, then Eq. 6 may be expanded by allowing *α* to be a function of temperature as determined by the Arrhenius equation,

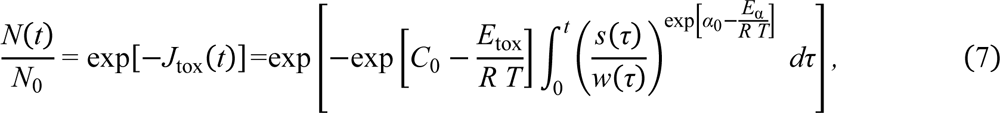

where *α*_0_ and *E*_α_ are fitted parameters. Equation 6 will be referred to as the Intracellular Molality Arrhenius Power (IMAP) model. It is possible the toxicity death rate associated with CPAs is not solely based on intracellular molality of CPAs. Indeed, cryoprotectants such as dimethyl sulfoxide (Me_2_SO) may create micro-pores in lipid membranes and change the permeability of ions across the membrane (52–54). If damage to the membrane is the key factor for loss of cell function, then the death rate may be better modelled as proportional to molality of CPA at the cell boundary. Using the basis of Eq. 7, and letting decay rate *r* be proportional to extracellular molality *m*_e_, an analytical solution may be taken when *m*_e_ remains constant such that,

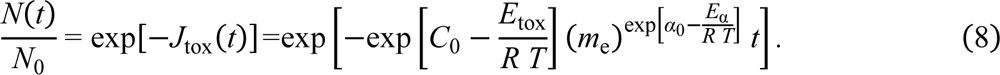

We present Eq. 8 as a novel model and it will be referred to as the Extracellular Molality Arrhenius Power (EMAP) model. However, during unloading or dehydration, intracellular molality of CPA may be more than extracellular molality. Equation 8 may be modified to take the maximum between intracellular CPA molality and extracellular CPA molality. Although not covered here, an alternative is to take the average intracellular and extracellular CPA molality.

#### Osmotic damage model

Taking a similar approach as above, osmotic damage may be derived using Eq. 4. We argue that osmotic damage decay rate is proportional to a power of volumetric change, ΔV, from resting state (taken as isotonic volume), taking the form,

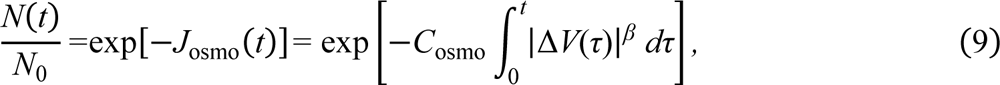

where Δ*V*(*t*)*=V*(*t*) − *V*_iso_, and *C*_osmo_ is a fitted constant. We present Eq. 9 as a novel model and it will be referred to as Time Dependent Osmotic Damage (TDOD) model. Note the absolute difference for change in volume (vertical bars) since we expect both swelling and shrinking to be harmful.

As many cells have different hypotonic compared to hypertonic osmotic limits, we will not assume *C*_osmo_ and *β* are the same for swelling and shrinking (55). Therefore, the constants must be fit independently to hypotonic and hypertonic challenge. Furthermore, it is possible that osmotic damage is also solution dependent (e.g. high concentrations of sucrose may interact differently with the membrane than NaCl).

The osmotic tolerance limit hypothesis predicts osmotic damage to be constant throughout time, whereas the TDOD model (Eq. 9) predicts osmotic damage to be time dependent. The time dependent osmotic damage model can use the 2P model to obtain dynamic volumetric change throughout time.

#### Temperature damage model

Temperature damage rate is a function of temperature difference, Δ*T*, (with reference to physiological temperature, *T*_phys_), integrating yields the form:

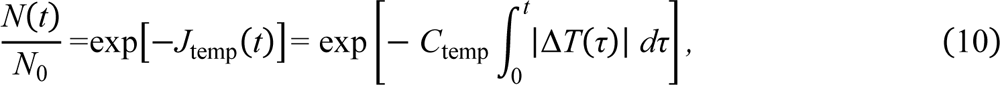

where Δ*T*(*t*)*=T*(*t*) − *T*_phys_, *T*(*t*) is average temperature of the cell at time *t*, while *C*_temp_ and *T*_phys_ are fitted constants. We note that absolute values are present due to the expectation that cells exposed to above, as well as below, physiological temperatures, will experience damage (and recognizing fitted constants may be different). Furthermore, many organisms may survive normally within a range of temperatures, therefore *T*_phys_ should be taken as the temperature at which temperature damage starts to occur (for respective cooling or heating). We present Eq. 10 as a novel model and it will be referred to as the Temperature and Time Dependent (TTD) model.

An alternative model asserts that chill injury is temperature dependent but not time dependent (assuming temperature equilibration between cell and environment). Integrating a first order approximation of population decay as a function of temperature yields,

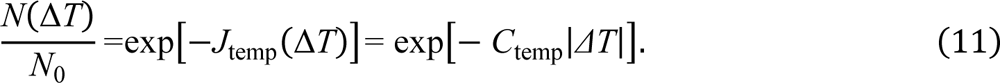

This equation assumes a single change in temperature and equilibration of the cells to that temperature. For dynamic changes in temperature, the largest change in temperature may be taken (i.e: | Δ*T |* = max[|*T*(*t*)−*T*_phys_ |], where *t* ranges from the start to the end of the protocol time and *T*(*t*) is average temperature of the cell at time *t*). We present Eq. 11 as a novel model and it will be referred to as the Temperature Dependent (TD) model.

#### Total damage model and optimization

The proportional survival of a population at the end of a cryopreservation protocol is related to the total damage, *J*_tot_, of that protocol. This total damage is asserted to be a linear combination of cytotoxicity, osmotic damage, and temperature damage such that, *J*_tot_=*J*_tox_+*J*_osmo_+*J*_temp_. This relation is directly derived by taking a first order approximation of population decay in the form,

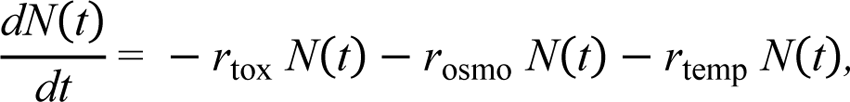

where *r*_tox_, *r*_osmo_, *r*_temp_, are the decay rates due to toxicity, osmotic damage and temperature, respectively. Solving gives,

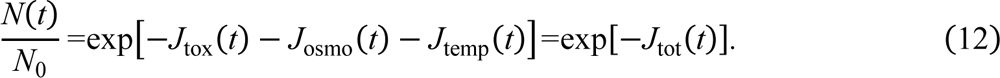

There are three types of continuous loading/unloading optimization approaches of CPA into and out of the cell: minimal time, minimal volume deviance, and minimal toxicity (14,36,56). For simplicity, we consider just loading protocols for a given goal intracellular CPA (in this case Me_2_SO) volume per solution volume (v/v), since if the cells cannot be successfully loaded to the goal CPA v/v then looking for an unloading protocol is not applicable. The first method is time optimization, whereby loading time is minimized by performing continuous loading while the cell is shrunken. Second, volume optimization whereby volumetric related damages are minimized by loading the cell close to or at isotonic volume. Lastly, cytotoxicity optimization, whereby cytotoxicity is minimized by loading the cell in a swollen state and quickly dehydrating the cell right before vitrification. Typically, these optimizations are performed in the context of osmotic tolerance limits; however, Eq. 12 enables an investigation of a spectrum between these approaches stated above. A search for an optimized loading protocol is the maximum survival as a function of the temperature during loading, the volume at which the cell is loaded, and the goal CPA v/v. Fig 1 depicts examples of minimal time, minimal volume deviance, and minimal toxicity during loading and the respective normalized survival throughout time given 30% goal CPA v/v%.

**Fig 1.**
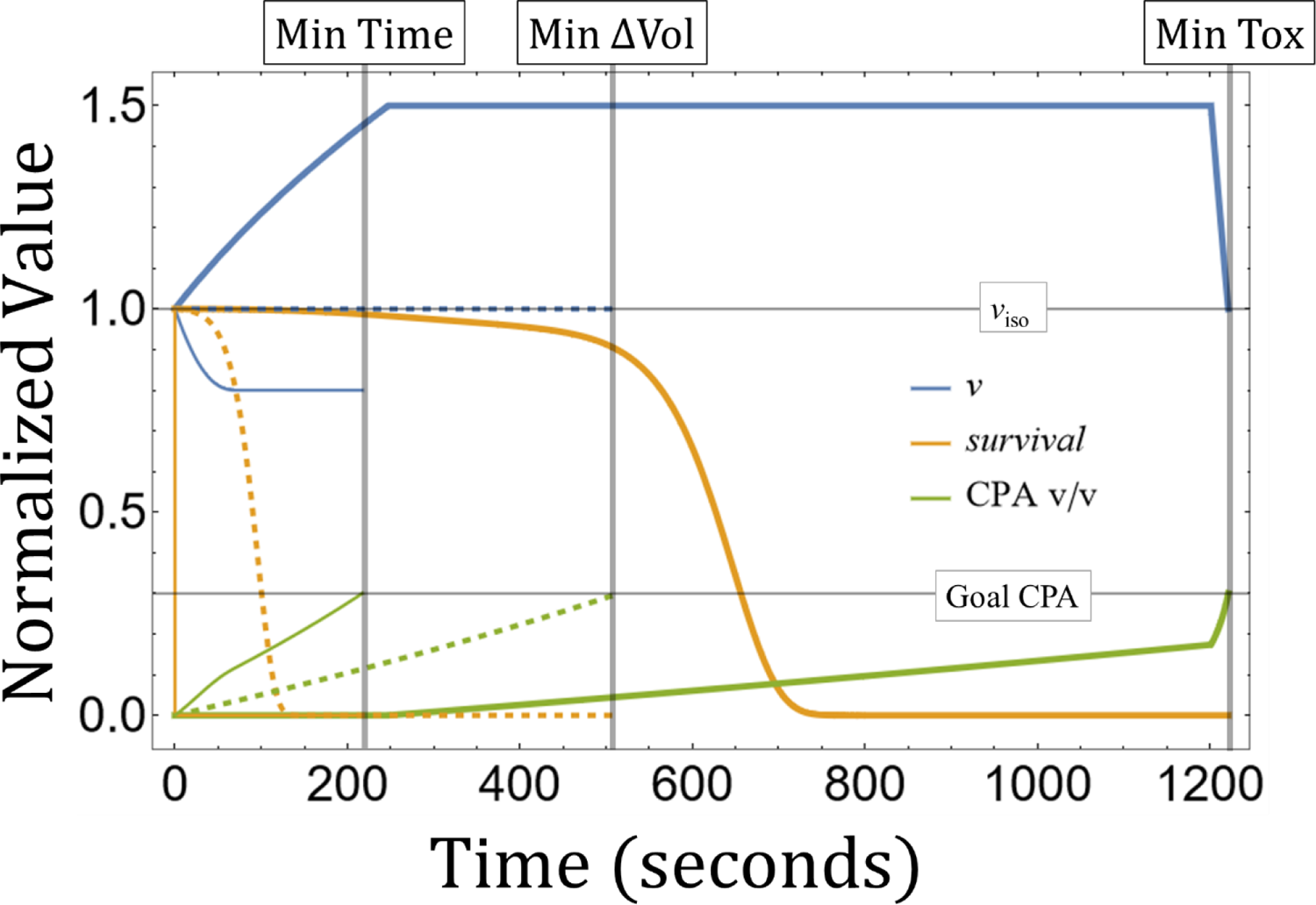
Example of optimized loading protocols for sea urchin (*P. lividus*) oocytes to 30% Me_2_SO v/v%. Solid thick lines represent minimal toxicity protocol, dotted lines represent minimal volume deviance protocol, and solid thin lines represent minimal time protocol. Transparent grey lines indicate the end of the respective loading protocol. Colors indicate relative volume (bleu), relative survival (orange), and intracellular CPA v/v (green).

### Gamete collection

Mature sea urchins (*Paracentrotus lividus*) were obtained from a conditioned broodstock from Toralla Marine Science Station, maintained for several months in a flow-through system under optimal conditions to enhance gonad maturation. Adults were dissected and gametes were collected directly from the gonads using a Pasteur pipette. A total of three females per experiment were used to collect oocytes and one male per experiment is used to reduce animal use. Gamete quality was checked before the experiment under light microscope, focusing on color and shape of oocytes, motility for sperm.

### Experimental Solutions and Reagents

Dimethyl sulfoxide, NaCl, and sucrose are from Sigma−Aldrich. Deionized (DI) water was obtained with a Vent Filter MPK01 from Milipore and seawater (SW) was filtered by 1 µm and UV sterilized. Where stated, solutions were created by adding the respective solute to seawater taken to be 1.0 osmol/kg. Hypotonic solutions were created by adding deionized water to seawater. Boyle van ‘t Hoff solutions were created by addition of NaCl with DI water.

### Measurement of Viability

Viability (cell survival) was measured using a functional assay (development to the 4-arm-pluteous stage). Following treatment, sea urchin oocytes were placed in 20 ml food−safe polypropylene vials with room temperature seawater and left for approximately 20 minutes to equilibrate before fertilization. Fertilization was done by addition sperm (15:1 sperm−oocyte ratio) into containers with densities of 20-40 oocytes/ml of seawater (57, 58). After 48 hours, development was arrested by addition of 40% buffered formalin. Survival was assessed by counting the number of individuals that develop to the 4-arm-pluteous stage (57, 58) over total count (at least 100 per replicate) with a Nikon Eclipse TE2000S microscope with the NIS elements software. For each experiment, survival was normalized with respect to the appropriate control.

### Boyle van ‘t Hoff (BVH) experimental design

NaCl solutions (486, 602, 778, 972, 1215, 1944, 3888 mOsm/kg) were prepared in Milli−Q water. Sea urchin oocytes were concentrated using a 40 µm mesh and deposited in Petri dishes containing NaCl solutions. Temperature was maintained at 18±1 °C throughout all the BvH treatments. Three replicates were assayed for each treatment. Oocyte/fertilized egg images were captured using SMZ 1500 binocular loupe and NIS Elements D software at 30 minute exposure. Digital images were analyzed and the major and minor semi−axes of each oocyte were measured. Each cell volume, *V*, was determined using the formula:

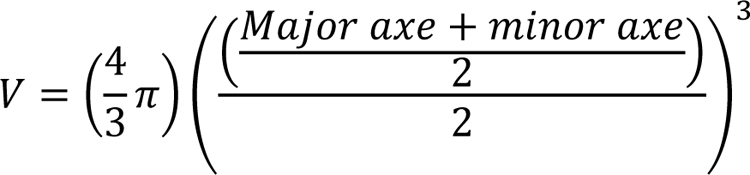

when the percentages of deviation between the mean and respective major and minor axes were less than the considered 10%. If this condition was not met, then the cell volume was calculated assuming prolate spheroid shape and using the following formula:

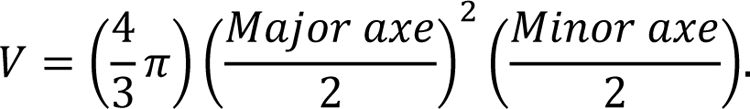

### Membrane characteristics experimental design

Sea urchin oocytes were exposed to seawater supplemented with 1.5 osmol/kg of Me_2_SO at room temperature (20±1 °C, with daily room temperatures logged explicitly and used in the analysis). Oocyte volume was recorded with an Nikon Eclipse TE2000S microscope for 22 minutes. At least 51 oocytes were recorded. Images were processed in ImageJ (59) and cross−sectional area of oocytes were recorded. The 2P model (Eq. 2) was fit to individual cell volumes throughout time. The Me_2_SO equilibration protocol was repeated at 10±1 °C and 6±1 °C and individual oocyte volumes were tracked throughout time. Temperatures were held constant in air-conditioned rooms. The Arrhenius equation for the 2P model was linearly fit by taking the log of the *L*_p_ and *P*_s_ values and plotting them against inverse temperature.

### Osmotic damage experimental design

To test the hypothesis of time-independent osmotic tolerance limits, oocytes were exposed to hypertonic solutions of seawater (UV filtered seawater at 1.0 osmol/kg) supplemented with 0.5, 1.0, and 1.5 osmol/kg of NaCl for duration periods of 2, 6, 15, and 30 minutes. Technician corrected exposure times were recorded for each treatment (e.g. 2.5 minutes instead of 2 minutes). To determine if hypertonic related damage is solution specific, the experiment was repeated with additions of sucrose in place of NaCl. Sucrose treatments included seawater (1.0 osmol/kg) supplemented with 0.5, 1.0, 1.5, and 2.0 osmol/kg sucrose for the same exposure times. Hypotonic damage was tested by diluting seawater by DI water exposing them to osmolalities of 0.8, 0.7, 0.6, and 0.5 osmol/kg for the same exposure periods. All experiments were done at room temperature (20 °C) and each treatment had three replicates each containing more than 600 oocytes per replicate (of which at least 100 were counted). All replicates were inside 20 ml food-safe polypropylene vials and solutions were filtered with a 20 µm polyethylene mesh. A control (three replicates) was kept in the same area throughout the duration of the experiment in seawater. Survival was calculated as described in Measurement of Viability section.

### Temperature damage experimental design

We tested effects of chill injury at three temperatures 20 °C, and 10 °C in air conditioned rooms (typical range of ±1 °C), and in an ice bath with an average temperature of approximately 1.7±2 °C (with observed temperature ranges from 0.4 to 3.0 °C and an SEM of ± 0.25 °C). Exposure times were 2, 6, 15, 30, 50, 75, and 90 minutes with three replicates (as described above). All treatments were conducted in seawater (1.0 osmol/kg). Survival was normalized with respect to the mean survival of oocytes throughout the 20 °C treatment. Survival was calculated as described in Measurement of Viability section.

### Cytotoxicity experimental design

We tested effects of cytotoxicity at three temperatures concurrently with the temperature damage experiment (see above). All treatments were conducted in seawater (1.0 osmol/kg) supplemented with 0.5 osmol/kg, 1.0 osmol/kg, or 1.5 osmol/kg Me_2_SO. Exposure times were 2, 6, 15, and 30 minutes with three replicates (as described above). Note, for ice bath treatment, only additions of 0.5 osmol/kg, and 1.0 osmol/kg Me_2_SO were performed. Eq. 9 was used to account for any osmotic damage during both loading and unloading of Me_2_SO. For each respective treatment, survival was normalized to the mean survival of oocytes at the respective temperature in seawater (1.0 osmol/kg). The cytotoxicity models and the osmotic damage model were informed by the 2P model (Eq. 2). Survival was calculated as described in Measurement of Viability section.

### Numerical optimization design

We used Equation 12 with respect to the verified damage models (Eq. 8, 9 and 11) to obtain relative population survival for the respective loading protocol. Continuous loading protocols may be defined by the temperature, loading volume, and goal CPA v/v. We varied temperature from 0 °C to 20 °C and varied relative loading volume from 0.5 to 3.0. The goal CPA v/v for vitrification of oocytes was checked from 5% to 50% CPA v/v% at intervals of 2%. We performed an exhaustive search within each respective domain.

### Analysis

All models, fits, and analysis were conducted in Mathematica (12.3v) (60). The BvH relation (Eq. 1) was fit through linear regression using Mathematica’s *LinearModelFit* function. A Durbin Watson score was obtained (first order autocorrelation test of residuals) with values close to zero indicating high autocorrelation and values close to 2 indicating no autocorrelation (61, 62).

Spearman Ranked correlation tests were performed for each chill damage treatment between temperature and time. All nonlinear models were fit to each respective dataset using the *NonLinearModelFit* function in Mathematica. A grid search was used to provide initial guesses for fitting parameters of Eq. 6-9. Comparison between models is conducted with goodness of fit with respect to the plotted data and respective model along with AIC and BIC. Adjusted R^2^ is presented for nonlinear models but conclusions must not be taken with respect to Adjusted R^2^ values alone (63). We note that we present R^2^ value for the Arrhenius equation, which, while linear, is a transformed dataset and therefore the presented R^2^ value must be viewed with this consideration. Replicates that were erroneously conducted were documented and removed from the analysis (e.g. failed to add semen at the respective time, or failed to effectively remove sucrose). Code and data may be found in the supplemental materials.

## Results

Equilibrated volumes of sea urchin oocytes were fit by the Boyle van ‘t Hoff relation (Fig 2A) with an R^2^ value of 0.704 (n=1103). The linear relation did not appear autocorrelated (Durbin Watson score of 2.32), indicating Eq. 1 is an appropriate model. The fitted osmotically active fraction, *w*_iso_, of *P. lividus* oocytes is 0.687±0.014 (SE), and the osmotically inactive fraction, *b*, is 0.311±0.017. We recorded exposed oocytes to 1.5 osmol/kg of Me_2_SO at 6 °C, 10 °C and 20 °C. We tracked individual oocytes throughout time and we fitted initial cell volume, *L*_p_ and *P*_s_ values (n=214). Fig 2B shows the fitted Arrhenius equation of log[*L*_p_] and log[*P*_s_] as a function of inverse temperature with values presented in Table 1. The activation energies *E*_lp_ and *E*_ps_ are 13.025±0.643 kcal/mol and 25.922±1.148 kcal/mol respectively. Fig 2C shows the mean of each normalized cell with respect to its initial volume and plotted is Eq. 2 using the average *L*_p_ and *P*_s_ values for the given temperature. While the 2P model fits well to the data, unexpectedly, oocytes exposed to Me_2_SO at 6 °C were larger on average than oocytes exposed at 10 °C.

**Fig 2.**
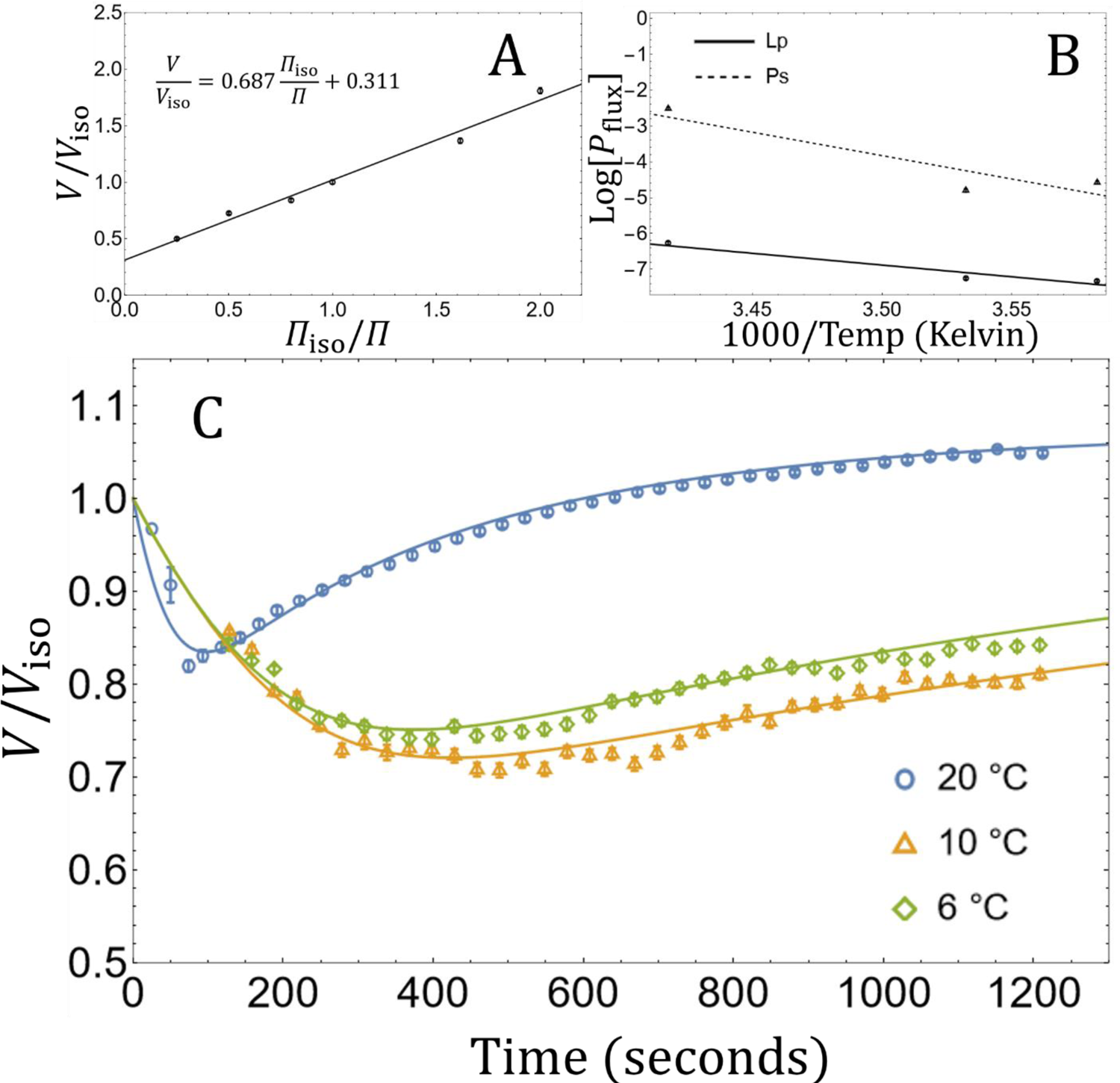
Osmotic characteristics of *P. lividus* oocytes. Symbols represent mean values and error bars represent SEM. Panel A, nondimensionalized Boyle van ‘t Hoff Plot (n=1103). Panel B, Arrhenius log plot where *P*_flux_ represents *L*_p_ and *P*_s_ (n=214). Panel C, normalized volume flux of oocytes in seawater (1.0 osmol/kg) with addition of 1.5 osmol/kg dimethyl sulfoxide at differing temperatures. Solid lines are the mean of the individually fitted 2P model with respect to each oocyte (n=214).<colcnt=1>

**Table 1.** Fitted flux parameters and Arrhenius equation fitted parameters (n=214). *E*_activation_ represents *E*_Lp_ and *E*_Ps_, while Arrhenius intercept represents *L*_0_ and *P*_0_

| Flux<br>Parameter | ($\mu\text{m atm}^{-1} \text{min}^{-1}$)<br>($\mu\text{m min}^{-1}$) | Arrhenius<br>intercept ± SE<br>(dimensionless) | $E_{\text{activation}} \pm$ SE<br>(mol/kcal) | $R^2$ |
| --- | --- | --- | --- | --- |
| $L_p$ | 0.118±0.005 @ 20 °C | 16.060±1.140 | 13.025±0.643 | 0.971 |
| | 0.044±0.001 @ 10 °C | | | |
| | 0.041±0.001 @ 6 °C | | | |
| $P_s$ | 5.036±0.198 @ 20 °C | 41.832±2.036 | 25.922±1.148 | 0.983 |
| | 0.541±0.030 @ 10 °C | | | |
| | 0.710±0.041 @ 6 °C | | | |

Using an *L*_p_ value in pure seawater of 0.191 μm atm^−1^ (9), the TDOD model (Eq. 9) was fit to seawater (1.00 osmol/kg) supplemented with NaCl (Fig 3A), with sucrose (Fig 3B) or diluted with DI water (Fig 3C). The fitted model appears to fit well in all cases (Fig 3) with high adjusted R^2^ values (adjR^2^>0.98) Noticeably, shrinking damages for NaCl and sucrose had a high fitted power, *β=*10.88±0.54, and *β=*11.29±0.61 respectively, while swelling damages for diluted seawater had a lower fitted power *β=*1.55± 0.17 (See Table 2). Furthermore, the parameter with respect to the NaCl treatment and the sucrose treatment are significantly different (parameter confidence region ellipsoids of Eq. 9 do not overlap with a confidence level of 95%). For clarity, fitted parameters are distinct between treatments sucrose and NaCl solutions. Thus, for a given time, and concentration, osmotic related damage to an oocyte in a sucrose solution is more damaging than NaCl solution (Fig 3A-B).

**Fig 3.**
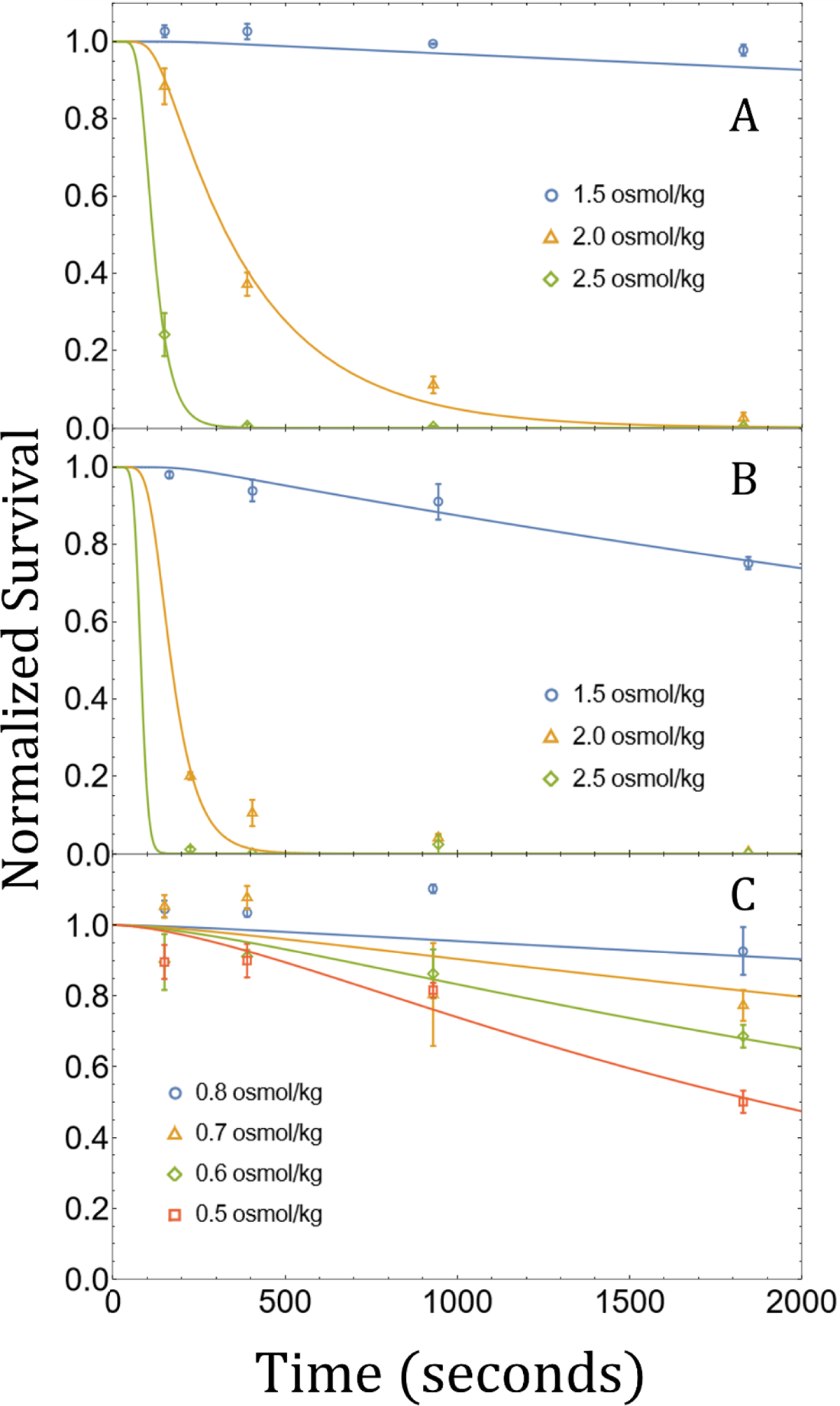
Time dependent osmotic damage model for multiple solutions. All plots are normalized to their respective controls (1.0 osmol/kg seawater). Listed osmolalities are total osmolalities of solution (e.g. 1.0 osmol/kg seawater + 0.5 osmol/kg NaCl). Symbols represent mean values and error bars represent SEM. Panel A, normalized survival of oocytes in NaCl + seawater. Panel B, normalized survival of oocytes in sucrose + seawater. Note 3.0 osmol/kg sucrose + seawater data are not plotted but all have 0% survival. Panel C, normalized survival of oocytes in Reverse Osmosis water + seawater.

**Table 2.** Fitted time dependent osmotic damage (TDOD) model for three different conditions.

| Solution | $C_{\text{osmo}} \pm \text{SE}$ | $\beta \pm \text{SE}$ | n | adjR<sup>2</sup> |
| --- | --- | --- | --- | --- |
| NaCl + SW | 6.55±2.55 | 10.88±0.54 | 35 | 0.995 |
| Sucrose + SW | 41.26±20.58 | 11.29±0.61 | 48 | 0.993 |
| DI + SW | 4.71±0.9 ×10<sup>-4</sup> | 1.55± 0.17 | 48 | 0.986 |

Temperature damage related to chill injury was investigated with respect to time and temperature dependent damage (TTD, Eq. 10), and solely temperature dependent damage (TD, Eq. 11). With respect to time and temperature, the fitted TTD model is shown in Fig 4A along with the fitted TD model, whereby they differ in estimated survival throughout time (TTD is dynamic with respect to time while TD is constant). TD is plotted with respect to temperature in Fig 4B. Performing a Spearman Rank correlation test for each temperature results in no significant correlation between time and cell survival (*p*>0.05; Table 3). While both models appear to have high adjusted R^2^ values (adjR^2^≥0.97), the TD model outperforms the TTD model in terms of adjR^2^, AIC, and BIC (Table 4).

**Fig 4.**
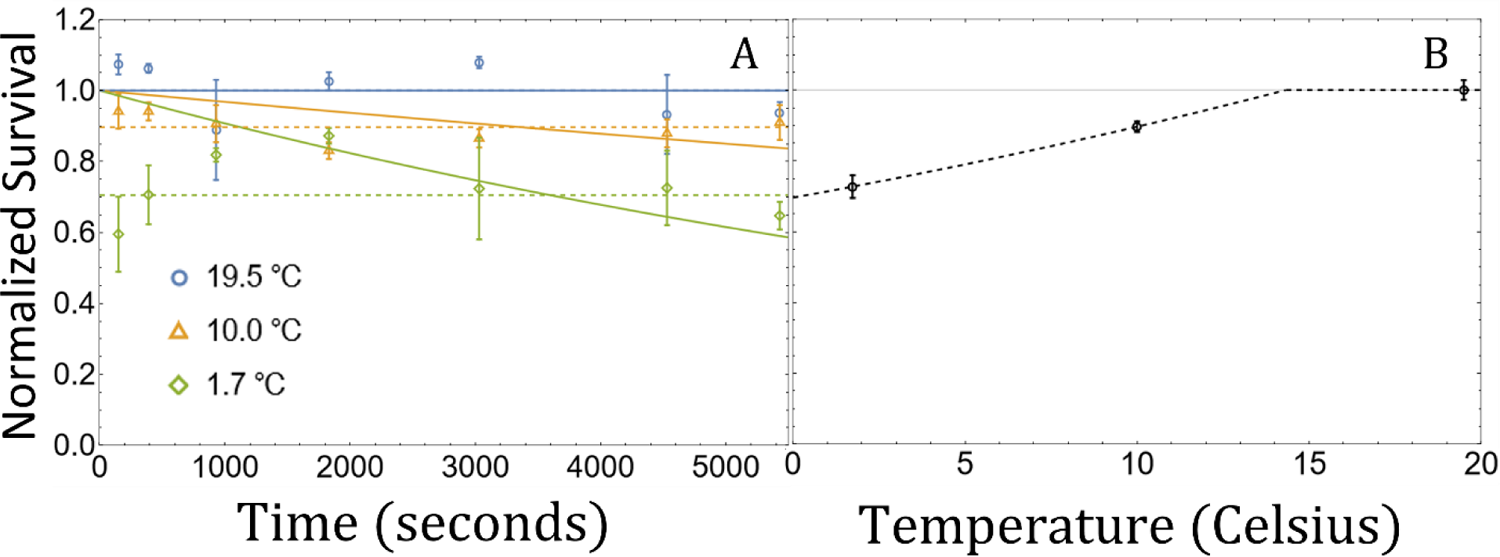
Temperature damage models of sea urchin oocytes. Data are normalized with respect to room temperature (19.5 °C) in 1.0 osmol/kg seawater. Ice bath had an average temperature of 1.7±2 °C. Symbols represent mean values and error bars represent SEM. Panel A, Normalized survival with respect to time (n=62). Solid lines represent the TTD model (Eq.10), while dotted lines represent the TD model (Eq. 11). Panel B, Normalized survival with respect to temperature (n=62). Dotted line represents the TD model (Eq. 11).

**Table 3.** Results of Spearman Rank correlation tests between time and survival at each respective temperature.

| Temperature (°C) | n | r-Statistic | *p* |
| --- | --- | --- | --- |
| 19.5 | 21 | −0.401 | 0.071 |
| 10 | 21 | −0.232 | 0.316 |
| 1.7 | 20 | −0.034 | 0.888 |

**Table 4.** Comparison of temperature damage models.

| Model | $C_{temp} \pm SE$ | $T_{phys} \pm SE$ | n | adjR<sup>2</sup> | AIC | BIC |
| --- | --- | --- | --- | --- | --- | --- |
| TTD<br>(Eq. 10) | 0.068±0.026 | 14.77±3.30 | 62 | 0.970 | −52.466 | −46.085 |
| TD<br>(Eq. 11) | 0.025±0.006 | 14.33±1.85 | 62 | 0.983 | −87.181 | −80.800 |

Using the fitted TDOD model (Eq. 9) to correct for any osmotic damage, the three cytotoxicity models (IMA, Eq. 6; IMAP, Eq. 7; and EMAP, Eq. 8) were fit and plotted across temperature and concentration (Fig 5). The EMAP model performed the best in terms of adjR^2^, AIC, and BIC when compared to IMA and IMAP (see Table 5 for respective values and fitted parameters). Furthermore, the EMAP model appears to best visually fit data across each temperature (Fig 5). Interestingly, the fitted power decreases for the EMAP model as temperature decreases, such that *α*=11.08 at 20 °C, *α*=5.51 at 10 °C, and *α*=2.96 at 1.7 °C, while the proportionality constant increases as temperature decreases. A similar trend is found for IMAP (i.e. power constant decreases but proportionality constant increases).

**Fig 5.**
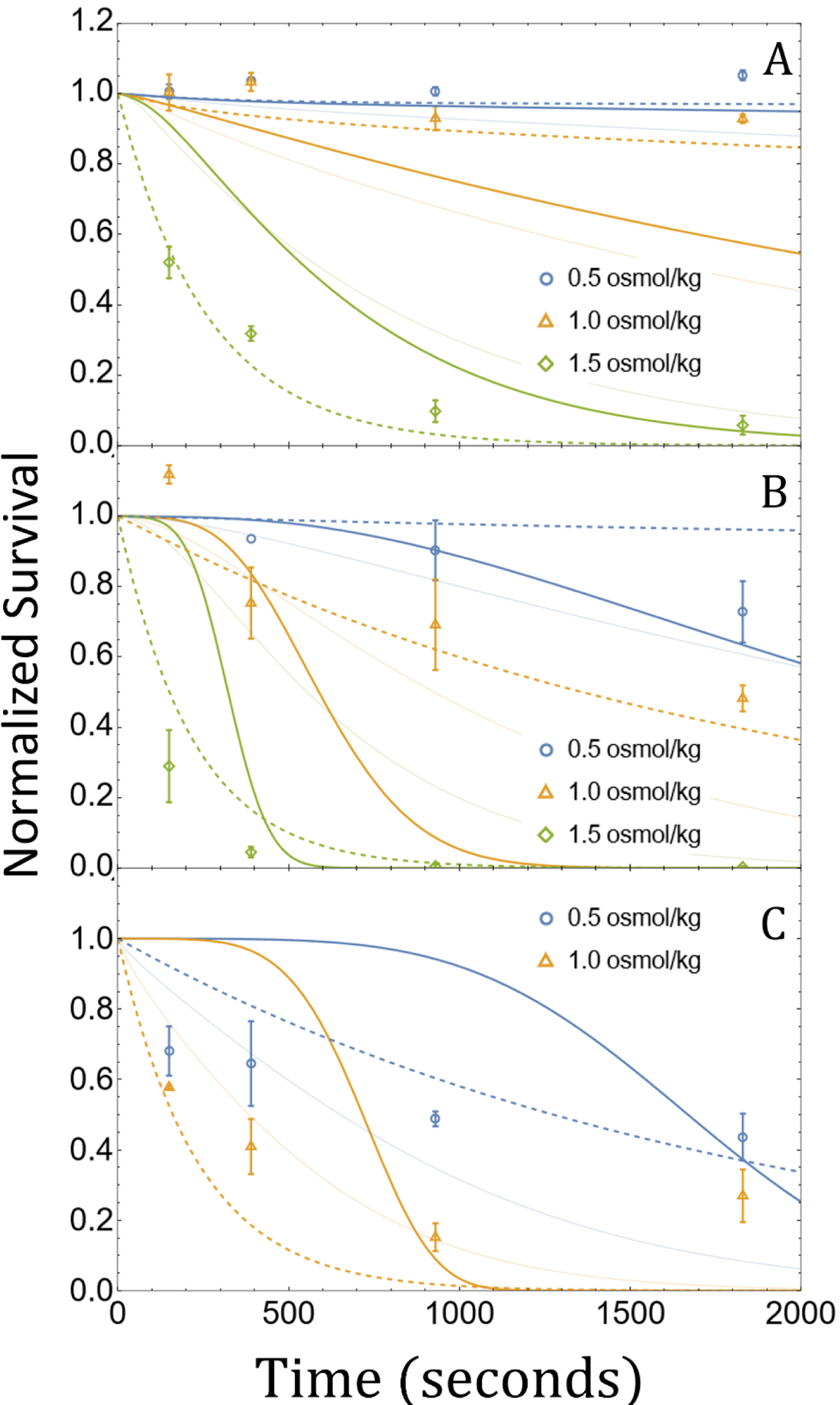
Cytotoxicity damage of dimethyl sulfoxide for sea urchin oocytes at different temperatures (n=89). Data are normalized to their respective control at their respective temperature. Listed osmolalities are the added osmolality of dimethyl sulfoxide to isotonic seawater (1.0 osmol/kg). Symbols represent mean values and error bars represent SEM. The solid thick line represents Eq. 6, the solid thin line represents Eq. 7, the dotted line represents Eq. 8. Panel A, normalized survival of oocytes with respect to time at room temperature (19.5 °C). Panel B, normalized survival of oocytes with respect to time in at 10 °C. Panel B, normalized survival of oocytes with respect to time in at 1.7±2 °C).

**Table 5.** Comparison of cytotoxicity models.

| Model | Fitted Parameters<br>(± SEM) | #Parameters | n | adjR<sup>2</sup> | AIC | BIC |
| --- | --- | --- | --- | --- | --- | --- |
| IMA<br>(Eq. 6) | $C_0 = -128.79 \pm 19.16$<br>$E_{tox} = -70.22 \pm 11.01$<br>$\alpha = 4.70 \pm 0.48$ | 3 | 89 | 0.813 | 44.480 | 54.435 |
| IMAP<br>(Eq. 7) | $C_0 = -44.88 \pm 11.06$<br>$E_{tox} = -21.55 \pm 6.34$<br>$\alpha_0 = 36.97 \pm 8.82$<br>$E_\alpha = 19.75 \pm 4.82$ | 4 | 89 | 0.848 | 27.176 | 39.619 |
| EMAP<br>(Eq. 8) | $C_0 = -80.55 \pm 8.62$<br>$E_{tox} = -41.01 \pm 4.77$<br>$\alpha_0 = 22.20 \pm 1.82$<br>$E_\alpha = 11.53 \pm 1.02$ | 4 | 89 | 0.953 | -78.119 | -65.675 |

Using osmotic damage defined by the TDOD model (Eq. 9), temperature damage defined by the TD model (Eq. 10), and cytotoxicity damage defined by the EMAP model (Eq. 8), we used the sum of these damages (Eq. 12) to perform an exhaustive grid search for the maximal survival during loading. Three examples of continuous loading are shown in Fig 1. We vary temperature (273 to 293 Kelvin) and relative loading volume (0.5 to 3.0) for the given goal CPA v/v (0.01 to 0.5), and obtain the respective survival at the end of loading. In Fig 6 we plot the optimal loading protocol with respect to the goal CPA (Me_2_SO) v/v (values above 0.31 not show). Sea urchin oocytes have approximately a 50% survival loading to about 13% Me_2_SO v/v%, while having a survival of 0.05% when loading to 25% Me_2_SO v/v%. Vitrification level solutions such as 40% and 50% Me_2_SO v/v%, have maximal survivals of 0%.

**Fig 6.**
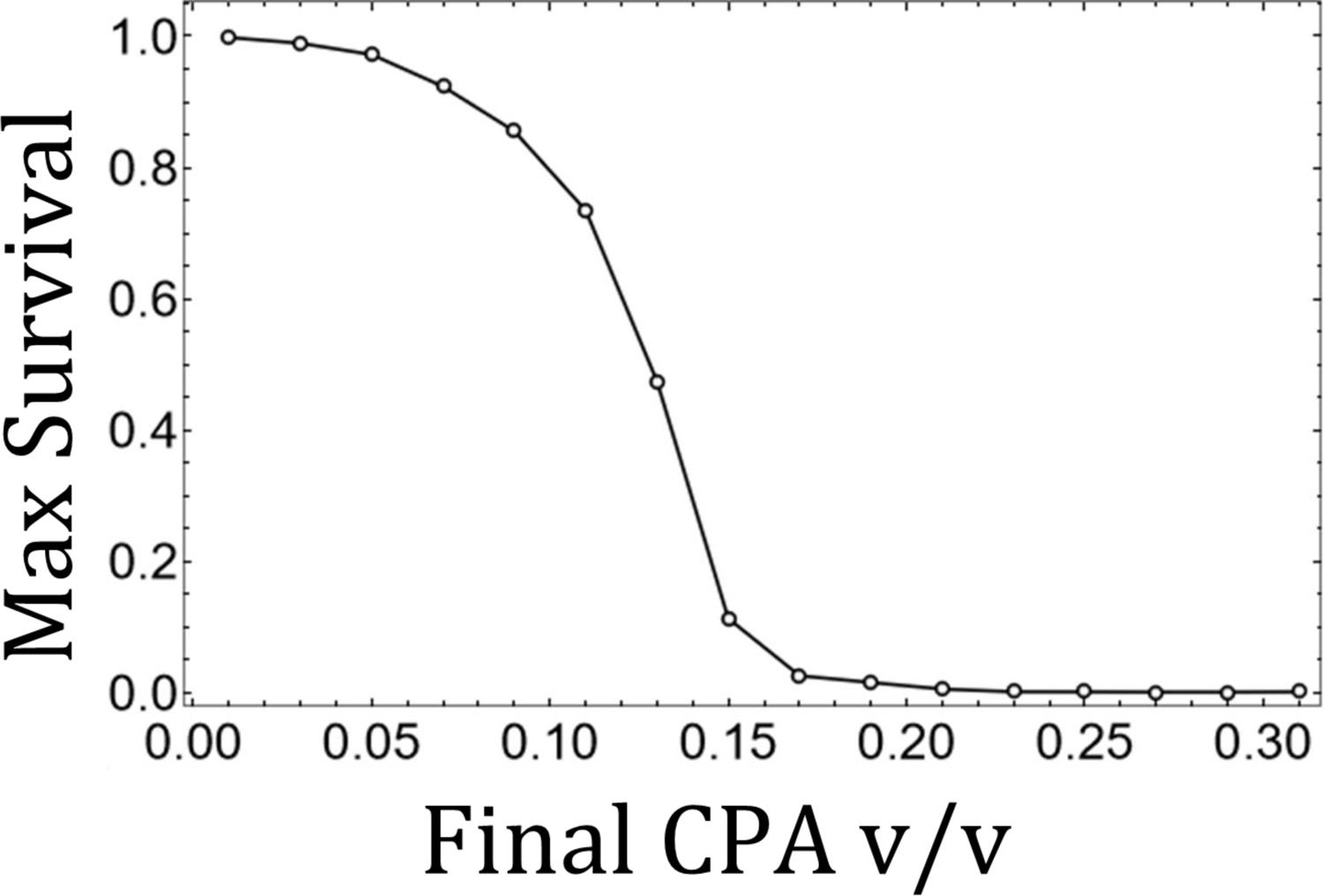
The maximal survival from an exhaustive grid search for optimal loading protocol. Max survival is plotted with respect to the final loaded CPA (Me_2_SO) v/v. Max survival values larger than 0.21 CPA v/v are < 0.001 (i.e. < 1% survival).

## DISCUSSION

### Cell and Membrane Osmotic Characteristics

Volume appears to be linearly related to inverse osmotic pressure as predicted for ideal osmometers following the BvH relation (Fig 2A). This supports the assumption that *Paracentrotus lividus* oocytes are ideal osmometers. Notably, the sea urchin *Evechinus chloroticus* oocyte were also found to be ideal osmometers (9).

We recorded many oocytes throughout their loading phase (n=214) and normalize these oocytes with respect to each individual fitted initial volume. Such a large sample size, and a correlated *L*_p_, *P*_s_, and *V*_iso_ parameters, provides a rich dataset. Unexpectedly, the mean population volume at 10 °C is lower than the mean population volume at 6 °C (Fig 2C). According to the Arrhenius equation, the mean volume at for the 10 °C treatment should be higher than the mean volume for volume at 6 °C treatment. A possible explanation may be due to differences between oocytes between treatments, as the treatments were done on separate days and different females the oocytes were collected from (3 females per day). However, an alternative explanation is that the log of permeability parameter is nonlinear with respect to inverse temperature (contrary to the Arrhenius equation). Lower temperatures induce membrane phase transition that may result in a change of membrane permeability (39,40,42), and hence, possibly explaining the increase in relative permeability of Me_2_SO with respect to water. Future work may investigate novel nonlinear permeability models and experimentally validate them with a larger array of temperatures.

### Osmotic Damage

The TDOD model fits osmotic damage throughout time with high accuracy (adjR^2^>0.96; Fig 3). Hypotonic solutions were less damaging than hypertonic solutions for the same volumetric deviance (swelling instead of shrinking). The high fitted powers of hypertonic solutions indicate sensitivity to shrinkage (Table 2). Furthermore, hypertonic relate damage was influenced by solution type: NaCl or Sucrose. Indeed, Davidson and colleagues (36) found that hypertonic sucrose solutions were more osmotically damaging for endothelial cells than NaCl solutions in general. The time dependent osmotic damage model suggests that both solution type and time spent under osmotic challenge is important for osmotic survival.

Time dependent osmotic damage has been found in several cell types (23,25,26). While osmotic tolerance limits may be a useful metric and are reported for many cell types, many of these reports do not investigate the time dependent nature of osmotic damage (18, 19). If the mechanism of osmotic damage is due to ultrastructural changes in the lipid membrane and cytoskeletal matrix, then time dependent osmotic damage may be a ubiquitous phenomenon across all cells. Indeed, using an osmotic tolerance limit with an equilibration time of 5 minutes may provide a poor approximation of cell survival over the course of an hour. Conversely, osmotic tolerance limits taken at 10 minutes may provide little predictive power in terms of cell survival on the order of seconds.

Future work investigating the mechanisms of osmotic damage may cover the role of membrane and cytoskeletal regulation. If damage is dependent on membrane and/or cytoskeletal changes throughout time, then we expect a decrease in temperature to reduce time dependent damage following an Arrhenius relation. It is possible that both the fitted constant, *C*_osmo_ and the fitted power, *β*, are related to temperature via the Arrhenius relation, as we found with cytotoxicity damage. Indeed, Wang and colleagues (26) found accumulated osmotic damage was temperature specific while Williams and Takahashi (64) found membrane lysis of sea urchin eggs was mitigated at lower temperatures during hypertonic challenge. Supplements that prevent membrane and cytoskeletal regulation may reduce time dependent osmotic damage. However, there may be a trade-off between time dependent osmotic damage and sensitivity to lysis during osmotic challenge, or loss of cell functionality with supplements such as cytochalasin-D or cytochalasin-B (27,29,65). Furthermore, some somatic cells such as MDCK cells have zero survival to hypotonic shock when treated with cytochalasin-D but have an increased survival when treated with phalloidin (which stabilizes f−actin) (66).

### Temperature Damage

For short time periods (tens of minutes) chill injury is best predicted with respect to environmental temperature and not time of exposure as described by the TD model (Eq. 11). The TTD model (Eq. 10) is built on the hypothesis that chill injury is both time and temperature dependent, however, we refuted this hypothesis on the time scale of tens of minutes since survival is not correlated to time at any temperature (Table 3). Notably the *p*-value of room temperature is 0.071, we argue this is a statistical coincidence since we do not expect oocyte aging to be influencing within an hour and a half in seawater since room temperature is not expected to be a damaging temperature.

Long term (several hours to days) temperature damage is known to be time dependent for sematic cells (39, 42) and for sea urchin (*Hemicentrotus pulcherrimus*) oocytes (43). While the mechanism of damage is poorly understood, it is known that a change in calcium permeability is the cause of chill injury for some somatic cells (39). At low temperatures lipid membranes may undergo a phase transition (40). This phase transition is accompanied by a change in permeability to ions such as calcium (present in seawater). Sea urchin oocytes are particularly sensitive to calcium permeability change as a small local flux of calcium may cause a cascading effect of opening voltage sensitive channels, releasing more calcium, resulting in a calcium wave (67). This calcium wave is a ubiquitous signal in oocytes and follows when the sperm head unites with the oocyte membrane (68). Subsequently, the oocyte may become nonresponsive to sperm, typically to avoid polyspermy (69, 70). However, if this calcium wave is caused by chilling, then oocytes may lose the ability to be properly fertilized and develop. Such an event would render the oocyte functionally dead.

### Cytotoxicity

Cytotoxicity is best described by considering CPA osmolality at the cell boundary, as captured by the EMAP model (Eq. 8). This model outperforms both the IMA model (Eq. 6), as well as the IMAP model (Eq. 7) which has the same number of parameters. The theory driving the IMA and IMAP models is that cytotoxicity is a metabolically driven phenomenon wherein CPAs interact with intracellular enzymes, proteins and organelles and hampers critical physiological processes. The EMAP model, on the other hand, only considers the rate of changes on the cell membrane, such that lysis, pore formation, and changes in ion permeabilities are correlated with extracellular CPA at the cell boundary (52, 54). Indeed, if the mechanism of chill injury for oocytes is a change in permeability to calcium and a subsequent calcium wave (and subsequent loss of functionality), then this same mechanism may be driving cytotoxicity. That is, damage to the membrane may result in a calcium wave and subsequent loss of functionality.

Interestingly, the power constant of cytotoxicity is negatively correlated with temperature (such that lower temperatures have lower power constants) whereas the proportionality constant is positively correlated with temperature (lower temperatures have higher proportionality constants). While the decrease in power indicates a reduction in sensitivity, the increase in proportionality indicates an overall increase in damage. For sea urchin oocytes, Me_2_SO is overall more damaging at lower temperatures than at physiological temperatures (Fig 4). This is contrary to expected if the mechanism of CPA damage is metabolically driven (21, 36). It is possible that at lower temperatures membrane pores created by Me_2_SO last longer, while repair to the sites in the lipid membrane is reduced due to phase transition and low temperatures. This mechanism may also explain the observed increase in *P*_s_ at 6 °C (Fig 2). Further, if these pores enlarge and stay enlarged for longer, then the likelihood of a calcium wave may increase, resulting in higher rates of a calcium wave (and thus the lowered survival rates observed). Further research may investigate these and other possible mechanisms.

### Optimization modelling

Cryopreserving sea urchin oocytes has yet to be reported. Our exhaustive search, covering the spectrum of minimal time, minimal volume change, and minimal cytotoxicity, shows that there is no current method of loading oocytes with enough CPA to obtain successful vitrification (14,36,56). High sensitivity to CPA and osmotic shock make the sea urchin oocyte very difficult to vitrify. Possible alternatives include liquidous tracking and optimized slow cooling, but it is unclear if oocytes cryopreserved with these methods will be recoverable.

On the other hand, reducing sensitivity to CPA and osmotic shock may be fruitful. Bovine oocyte cryopreservation is improved with the use of calcium chelators and ion channel blockers (71, 72). These additions reduce the likelihood of a calcium wave from initiating (68,69,73). Blocking loss of functionality may be a key step in successful cryopreservation of sea urchin oocytes.

The determination of the appropriate equations for *J*_tot_ may be used for other cells and enable a more accurate and robust search for cryopreservation protocols. These models provide insight into damages and help pinpoint mechanisms preventing cryopreservation. For instance, some cell types may not be chill sensitive, while others may have cytotoxicity better described by intracellular molality. Furthermore, we argue that the utility of constant osmotic tolerance limits must be tested in light of a trade-off between time dependent osmotic damage and cytotoxicity during loading and unloading of CPA.

## Conclusion

We provide a novel model that rationally combines volumetric effects, solution effects, and temperature effects to predict total population survival throughout time. Traditional approaches to modelling osmotic damage, cytotoxic, and chill injury have been found to be inadequate or non-existent in the case of sea urchin oocytes. Our novel models provide a robust approximation of cell survival across time, temperature, and osmotic conditions, and highlight important mechanisms of damages that are often left uninvestigated. Future work may pinpoint these mechanisms of damages, providing insight into cellular stresses, adaptations, and responses.

## Acknowledgments

We graciously thank the excellent review and editing support provided by Robyn Shuttleworth. We graciously thank the aid by Sara Campos in performing osmotic damage experiments.

## Notes

### Competing Interest Statement

The authors have declared no competing interest.

## References

1. Hagen NT. Echinoculture: from fishery enhancement to closed cycle cultivation. World Aquac 275–19. 1996;27 (4):6–19.

2. Hose JE. Potential uses of sea urchin embryos for identifying toxic chemicals: Description of a bioassay incorporating cytologic, cytogenetic and embryologic endpoints. J Appl Toxicol. 1985;5(4):245–54.

3. Agnello M, Roccheri MC. Apoptosis: Focus on sea urchin development. Vol. 15, Apoptosis. 2010. p. 322–30.

4. Hogan B. Sea urchin development. Nature. 1974;247(5437):166–166.

5. Rezazadeh Valojerdi M, Eftekhari-Yazdi P, Karimian L, Hassani F, Movaghar B. Vitrification versus slow freezing gives excellent survival, post warming embryo morphology and pregnancy outcomes for human cleaved embryos. J Assist Reprod Genet. 2009;26(6):347–54.

6. Gasparrini B, Attanasio L, De Rosa A, Monaco E, Di Palo R, Campanile G. Cryopreservation of in vitro matured buffalo (Bubalus bubalis) oocytes by minimum volumes vitrification methods. Anim Reprod Sci. 2007;98(3–4):335–42.

7. Arav A. Cryopreservation of oocytes and embryos. Vol. 81, Theriogenology. 2014. p. 96– 102.

8. Zhou XL, Al Naib A, Sun DW, Lonergan P. Bovine oocyte vitrification using the Cryotop method: Effect of cumulus cells and vitrification protocol on survival and subsequent development. Cryobiology. 2010;61(1):66–72.

9. Adams SL, Kleinhans FW, Mladenov P V., Hessian PA. Membrane permeability characteristics and osmotic tolerance limits of sea urchin (Evechinus chloroticus) eggs. Cryobiology. 2003;47(1):1–13.

10. Asahina E, Takahashi T. Cryopreservation of sea urchin eggs. Cryobiology. 1978;15(6):688–9.

11. Paredes E. Biobanking of a marine invertebrate model organism: The sea urchin. Vol. 4, Journal of Marine Science and Engineering. 2016.

12. Pegg DE. Principles of cryopreservation. In: Preservation of Human Oocytes: From Cryobiology Science to Clinical Applications. 2009. p. 12–24.

13. Fahy GM, Wowk B. Principles of cryopreservation by vitrification. Methods Mol Biol. 2015;1257:21–82.

14. Benson JD. Chapter 3 modeling and optimization of cryopreservation. In: Methods in Molecular Biology [Internet]. Springer, New York, NY; 2015 [cited 2018 Dec 17]. p. 83–119. Available from: http://link.springer.com/10.1007/978-1-4939-2193-5_3

15. Anderson DM, Benson JD, Kearsley AJ. Foundations of modeling in cryobiology—III: Inward solidification of a ternary solution towards a permeable spherical cell in the dilute limit. Cryobiology. 2020;92:34–46.

16. Nobel PS. The Boyle-Van’t Hoff relation. J Theor Biol. 1969;23(3):375–9.

17. Van’t Hoff J. The function of osmotic pressure in the analogy between solutions and gases. Proc Phys Soc London. 1887;9(1):307–34.

18. Blässe AK, Oldenhof H, Ekhlasi-Hundrieser M, Wolkers WF, Sieme H, Bollwein H. Osmotic tolerance and intracellular ion concentrations of bovine sperm are affected by cryopreservation. Theriogenology [Internet]. 2012 Oct 1 [cited 2018 May 22];78(6):1312–20. Available from: http://www.ncbi.nlm.nih.gov/pubmed/22819283

19. Kashuba Benson CM, Benson JD, Critser JK. An improved cryopreservation method for a mouse embryonic stem cell line. Cryobiology [Internet]. 2008 Apr 1 [cited 2018 May 22];56(2):120–30. Available from: https://www.sciencedirect.com/science/article/pii/S0011224007003495

20. Lawson A, Mukherjee IN, Sambanis A. Mathematical modeling of cryoprotectant addition and removal for the cryopreservation of engineered or natural tissues. Cryobiology. 2012;64(1):1–11.

21. Benson JD, Kearsley AJ, Higgins AZ. Mathematical optimization of procedures for cryoprotectant equilibration using a toxicity cost function. Cryobiology [Internet]. 2012 Jun 1 [cited 2019 Apr 1];64(3):144–51. Available from: https://www.sciencedirect.com/science/article/pii/S0011224012000028

22. Davidson AF, Benson JD, Higgins AZ. Mathematically optimized cryoprotectant equilibration procedures for cryopreservation of human oocytes. Theor Biol Med Model. 2014;11(1).

23. Zawlodzka S, Takamatsu H. Osmotic injury of PC-3 cells by hypertonic NaCl solutions at temperatures above 0°C. Cryobiology [Internet]. 2005 Feb 1 [cited 2018 Dec 17];50(1):58–70. Available from: https://www.sciencedirect.com/science/article/pii/S0011224004001385

24. Morris CE, Homann U. Cell surface area regulation and membrane tension. J Membr Biol [Internet]. 2001 Jan [cited 2018 Dec 17];179(2):79–102. Available from: http://link.springer.com/10.1007/s002320010040

25. Liu J, Mullen S, Meng Q, Critser J, Dinnyes A. Determination of oocyte membrane permeability coefficients and their application to cryopreservation in a rabbit model. Cryobiology. 2009;59(2):127–34.

26. Wang L, Liu J, Zhou G Bin, Hou YP, Li JJ, Zhu SE. Quantitative investigations on the effects of exposure durations to the combined cryoprotective agents on mouse oocyte vitrification procedures. Biol Reprod. 2011;85(5):884–94.

27. Wang CL, Xu HY, Xie L, Lu YQ, Yang XG, Lu SS, et al. Stability of the cytoskeleton of matured buffalo oocytes pretreated with cytochalasin B prior to vitrification. Cryobiology. 2016;72(3):274–82.

28. Horvath G, Seidel GE. Vitrification of bovine oocytes after treatment with cholesterol-loaded methyl-β-cyclodextrin. Theriogenology. 2006;66(4):1026–33.

29. Fujihira T, Kishida R, Fukui Y. Developmental capacity of vitrified immature porcine oocytes following ICSI: Effects of cytochalasin B and cryoprotectants. Cryobiology. 2004;49(3):286–90.

30. Sheetz MP. Cell control by membrane-cytoskeleton adhesion. Nat Rev Mol Cell Biol [Internet]. 2001 May 1 [cited 2019 Apr 26];2(5):392–6. Available from: http://www.nature.com/articles/35073095

31. Taglieri DM, Delfín DA, Monasky MM. Cholesterol regulation of PIP2: Why cell type is so important. Front Physiol [Internet]. 2013 [cited 2019 May 3];3 JAN:492. Available from: http://www.ncbi.nlm.nih.gov/pubmed/23316171

32. Senju Y, Kalimeri M, Koskela E V., Somerharju P, Zhao H, Vattulainen I, et al. Mechanistic principles underlying regulation of the actin cytoskeleton by phosphoinositides. Proc Natl Acad Sci U S A. 2017;114(43):E8977–86.

33. Sun M, Northup N, Marga F, Huber T, Byfield FJ, Levitan I, et al. The effect of cellular cholesterol on membrane-cytoskeleton adhesion. J Cell Sci. 2007;120(13):2223–31.

34. Fahy GM, Wowk B, Wu J, Paynter S. Improved vitrification solutions based on the predictability of vitrification solution toxicity. Cryobiology. 2004;48(1):22–35.

35. Best BP. Cryoprotectant Toxicity: Facts, Issues, and Questions. Rejuvenation Res. 2015;18(5):422–36.

36. Davidson AF, Glasscock C, McClanahan DR, Benson JD, Higgins AZ. Toxicity Minimized Cryoprotectant Addition and Removal Procedures for Adherent Endothelial Cells. PLoS One. 2015;10(11).

37. Wang X, Hua TC, Sun DW, Liu B, Yang G, Cao Y. Cryopreservation of tissue-engineered dermal replacement in Me2SO: Toxicity study and effects of concentration and cooling rates on cell viability. Cryobiology. 2007;55(1):60–5.

38. Elmoazzen HY, Poovadan A, Law GK, Elliott JAW, McGann LE, Jomha NM. Dimethyl sulfoxide toxicity kinetics in intact articular cartilage. Cell Tissue Bank. 2007;8(2):125– 33.

39. Bayley JS, Winther CB, Andersen MK, Grønkjær C, Nielsen OB, Pedersen TH, et al. Cold exposure causes cell death by depolarizationmediated Ca2+ overload in a chill-susceptible insect. Proc Natl Acad Sci U S A. 2018;115(41):E9737–44.

40. Quinn PJ. A lipid-phase separation model of low-temperature damage to biological membranes. Cryobiology. 1985;22(2):128–46.

41. Overgaard J, Macmillan HA. The Integrative Physiology of Insect Chill Tolerance. Vol. 79, Annual Review of Physiology. 2017. p. 187–208.

42. Nedvěd O, Lavy D, Verhoef HA. Modelling the time-temperature relationship in cold injury and effect of high-temperature interruptions on survival in a chill-sensitive collembolan. Funct Ecol. 1998;12(5):816–24.

43. Ohyama Y, Asahina É. Supercooling injury in the egg cell of the sea urchin. Cryobiology. 1972;9(1).

44. MacMillan HA, Sinclair BJ. The role of the gut in insect chilling injury: Cold-Induced disruption of osmoregulation in the fall field cricket, gryllus pennsylvanicus. J Exp Biol. 2011;214(5):726–34.

45. Koštál V, Yanagimoto M, Bastl J. Chilling-injury and disturbance of ion homeostasis in the coxal muscle of the tropical cockroach (Nauphoeta cinerea). Comp Biochem Physiol - B Biochem Mol Biol. 2006;143(2):171–9.

46. Koštál V, Vambera J, Bastl J. On the nature of pre-freeze mortality in insects: Water balance, ion homeostasis and energy charge in the adults of Pyrrhocoris apterus. J Exp Biol. 2004;207(9):1509–21.

47. Kay AG, Hoyland JA, Rooney P, Kearney JN, Pegg DE. A liquidus tracking approach to the cryopreservation of human cartilage allografts. Cryobiology. 2015;71(1).

48. Benson JD. Some comments on recent discussion of the Boyle van’t Hoff relationship. Cryobiology. 2012;64(2):118–20.

49. Katkov II. A two-parameter model of cell membrane permeability for multisolute systems. Cryobiology. 2000;40(1):64–83.

50. Benson JD, Chicone CC, Critser JK. Exact solutions of a two parameter flux model and cryobiological applications. Cryobiology. 2005;50(3):308–16.

51. Jacobs MH, Stewart DR. A simple method for the quantitative measurement of cell permeability. J Cell Comp Physiol. 1932;1(1):71–82.

52. Fernández ML, Reigada R. Effects of dimethyl sulfoxide on lipid membrane electroporation. J Phys Chem B. 2014;118(31).

53. Gurtovenko AA, Anwar J. Modulating the structure and properties of cell membranes: The molecular mechanism of action of dimethyl sulfoxide. J Phys Chem B. 2007;111(35):10453–60.

54. He F, Liu W, Zheng S, Zhou L, Ye B, Qi Z. Ion transport through dimethyl sulfoxide (DMSO) induced transient water pores in cell membranes. Mol Membr Biol. 2012;29(3– 4):107–13.

55. Kashuba CM, Benson JD, Critser JK. Rationally optimized cryopreservation of multiple mouse embryonic stem cell lines: I-Comparative fundamental cryobiology of multiple mouse embryonic stem cell lines and the implications for embryonic stem cell cryopreservation protocols. Cryobiology [Internet]. 2014 Apr 1 [cited 2018 Jun 6];68(2):166–75. Available from: http://www.ncbi.nlm.nih.gov/pubmed/24384367

56. Benson JD, Higgins AZ, Desai K, Eroglu A. A toxicity cost function approach to optimal CPA equilibration in tissues. Cryobiology. 2018 Feb 1;80:144–55.

57. Paredes E, Bellas J, Costas D. Sea urchin (Paracentrotus lividus) larval rearing - Culture from cryopreserved embryos. Aquaculture. 2015;437:366–9.

58. Saco-Álvarez L, Durán I, Ignacio Lorenzo J, Beiras R. Methodological basis for the optimization of a marine sea-urchin embryo test (SET) for the ecological assessment of coastal water quality. Ecotoxicol Environ Saf. 2010;73(4):491–9.

59. Schindelin J, Arganda-Carreras I, Frise E, Kaynig V, Longair M, Pietzsch T, et al. Fiji: An open-source platform for biological-image analysis. Vol. 9, Nature Methods. 2012.

60. Wolfram Research Inc. Mathematica. Champaign, Illinois: Wolfram Research, Inc.; 2018.

61. Durbin J, Watson GS. Testing for Serial Correlation in Least Squares Regression: I. Biometrika. 1950 Dec;37(3/4):409.

62. Ali MM. Durbin-watson and generalized durbin-watson tests for autocorrelations and randomness. J Bus Econ Stat. 1987;5(2):195–203.

63. Spiess AN, Neumeyer N. An evaluation of R2as an inadequate measure for nonlinear models in pharmacological and biochemical research: A Monte Carlo approach. BMC Pharmacol. 2010;10.

64. Williams RJ, Takahashi T. The role of osmotic stress and surface energy in freezing injury of sea urchin eggs. Comp Biochem Physiol -- Part A Physiol. 1982;73(4):621–6.

65. Takamatsu H, Takeya R, Naito S, Sumimoto H. On the mechanism of cell lysis by deformation. J Biomech [Internet]. 2005 Jan 1 [cited 2019 Jul 11];38(1):117–24. Available from: https://www.sciencedirect.com/science/article/pii/S0021929004001356?via%3Dihub

66. Clegg JS. MDCK Cells Under Severely Hypoosmotic Conditions. In: Mechanics of Swelling. 1992. p. 267–92.

67. Shen SS, Buck WR. Sources of calcium in sea urchin eggs during the fertilization response. Dev Biol. 1993;157(1):157–69.

68. Whitaker M. Calcium at fertilization and in early development. Vol. 86, Physiological Reviews. 2006.

69. Gardner AJ, Evans JP. Mammalian membrane block to polyspermy: new insights into how mammalian eggs prevent fertilisation by multiple sperm. Reprod Fertil Dev [Internet]. 2006;18:53–61. Available from: www.publish.csiro.au/journals/rfd

70. Cheeseman LP, Boulanger J, Bond LM, Schuh M. Two pathways regulate cortical granule translocation to prevent polyspermy in mouse oocytes. Nat Commun. 2016;7.

71. Wang N, Hao HS, Li CY, Zhao YH, Wang HY, Yan CL, et al. Calcium ion regulation by BAPTA-AM and ruthenium red improved the fertilisation capacity and developmental ability of vitrified bovine oocytes. Sci Rep. 2017;7(1).

72. Sanaei B, Movaghar B, Valojerdi MR, Ebrahimi B, Bazrgar M, Jafarpour F, et al. An improved method for vitrification of in vitro matured ovine oocytes; beneficial effects of Ethylene Glycol Tetraacetic acid, an intracellular calcium chelator. Cryobiology. 2018 Oct 1;84:82–90.

73. Wozniak KL, Carlson AE. Ion channels and signaling pathways used in the fast polyspermy block. Mol Reprod Dev [Internet]. 2020 Mar 1 [cited 2022 Jun 10];87(3):350–7. Available from: https://onlinelibrary.wiley.com/doi/full/10.1002/mrd.23168

